# Early exposure to UV radiation causes telomere shortening and poorer condition later in life

**DOI:** 10.1101/2021.12.21.473748

**Authors:** Niclas U. Lundsgaard, Rebecca L. Cramp, Craig E. Franklin

## Abstract

Determining the contribution of elevated ultraviolet-B radiation (UVBR; 280 – 315 nm) to amphibian population declines is being hindered by a lack of knowledge about how different acute UVBR exposure regimes during early life history stages might affect post-metamorphic stages via long-term carryover effects. We acutely exposed tadpoles of the Australian green tree frog (*Litoria caerulea*) to a combination of different UVBR irradiances and doses in a multi-factorial experiment, and then reared them to metamorphosis in the absence of UVBR to assess carryover effects in subsequent juvenile frogs. Dose and irradiance of acute UVBR exposure influenced carryover effects into metamorphosis in somewhat opposing manners. Higher doses of UVBR exposure in larvae yielded improved rates of metamorphosis. However, exposure at a high irradiance resulted in frogs metamorphosing smaller in size and in poorer condition than frogs exposed to low and medium irradiance UVBR as larvae. We also demonstrate some of the first empirical evidence of UVBR-induced telomere shortening *in vivo*, which is one possible mechanism for life-history trade-offs impacting condition post-metamorphosis. These findings contribute to our understanding of how acute UVBR exposure regimes in early life affect later life-history stages, which has implications for how this stressor may shape population dynamics.

**SUMMARY STATEMENT:** We demonstrate physiological carryover effects in amphibians that link larval UV exposure to detrimental impacts on juvenile frogs, including telomere shortening, which has implications for how UV shapes amphibian populations.

## INTRODUCTION

Solar ultraviolet-B radiation (UVBR; 280 to 315 nm) is a high-energy electromagnetic radiation and a pervasive stressor for many organisms (Williamson et al. 2019). In addition to facilitating endogenous vitamin D_3_ synthesis (Stiffler 1993; Antwis and Browne 2009), this genotoxic stressor can also form pyrimidine dimer lesions in DNA that disrupt transcription and replication, which can in turn lead to cancer, cell apoptosis and tissue damage (Mitchell and Nairn 1989; Batista et al. 2009; Londero et al. 2019). UVBR also causes oxidative stress through the production of reactive oxygen species (ROS; Heck et al. 2003; Kazerouni et al. 2016), although ultraviolet-A radiation (UVAR; 315-400 nm) is more potent in this regard (Schuch et al. 2017). Despite organisms having DNA repair mechanisms to remove UVBR-induced DNA damage (Sancar and Sancar 1988; Sancar and Tang 1993; Schuch et al., 2015b), even minor increases in UVBR irradiance may be sufficient to tip DNA damage rates beyond the capacity for repair, which can have detrimental downstream effects on organismal condition and survival (Blaustein et al. 1994, Pandelova et al. 2006, Schuch et al. 2015b).

Stratospheric ozone depletion and changes in climate have caused widespread increases in the irradiance and fluctuation of UVBR in regions of amphibian decline (Farman et al. 1985; Kerr and McElroy 1993; Herman et al. 1996; McKenzie et al. 1999; Middleton et al. 2001; Schuch et al. 2015a). Ambient UVBR can have a range of lethal and sublethal effects across many amphibian species (Blaustein et al. 1995; Tietge et al. 2001; Blaustein et al. 2005; Bancroft et al. 2008; Croteau et al. 2008; Cramp and Franklin 2018), and these effects may be exacerbated by interactions with other factors including disease, pollutants and temperature (Blaustein and Keisecker 2002; Blaustein et al. 2003; Alton and Franklin 2017; Lundsgaard et al. 2020). However, two significant gaps in this body of literature hinder the effective extrapolation of these individual-level effects of UVBR to population-scale effects and declines. Firstly, it is not known which parameters of UVBR exposure determine amphibian health risk, be it irradiance, dose or exposure duration (Lundsgaard et al. 2021), which leads to discrepancies in the estimated risks posed to amphibians by field measurements of UVBR (Palen et al. 2002; Blaustein et al. 2004; Olker et al. 2013). Secondly, most research has focussed on early life-history stages which are most susceptible to UVBR exposure, but that do not strongly drive population dynamics (Vonesh and De la Cruz 2002; Vonesh 2005).

Post-metamorphic life-history stages have the strongest influence on amphibian population dynamics (Biek et al. 2002) but are typically overlooked in UVBR research due to their mainly nocturnal habits (Bancroft et al. 2008). What is rarely considered are potential physiological carryover effects that link embryonic and larval UVBR exposure to impacts on post-metamorphic life stages, despite the implications that such effects would have for amphibian population dynamics (Van Allen et al. 2010).

Carryover effects are consequences of an early development experience that persist for a time after the experience that causes them ceases, such that cause and effect are separated by a measurable transitional period (O’Connor et al. 2014). For example, environmental factors encountered by embryonic and larval amphibians, including contaminants, predation, aquatic pH, food availability and conspecific density, have been shown to affect traits including locomotion, morphology, growth and survival in later life-history stages (Merila et al. 2000; Relyea 2001; Rasanen et al. 2002; Vonesh 2005; Chelgren et al. 2006; Tejedo et al. 2010; Van Allen et al. 2010; Boes and Benard 2013; Touchon et al. 2013; Rumrill et al. 2018).

Only a few studies have explicitly tested carryover effects of UVBR exposure in amphibians, with detrimental effects of exposure during embryonic and larval stages oftentimes only manifesting in later life-history stages (Smith et al. 2000; Pahkala et al. 2001, 2003; Belden and Blaustein 2002; Ceccato et al. 2016). Such ‘latent effects’ are becoming increasingly apparent across taxa (Pechenik 2006), yet little is known about the mechanistic basis for these effects, with changes in energy balance, oxidative stress, epigenetic modifications, and telomere lengths all implicated (O’Connor et al. 2014; Young 2018).

Telomeres are non-coding DNA sequence repeats (TTAGGG in vertebrates) on the ends of chromosomes (in eukaryotes) and serve a protective role in maintaining genome stability (O’Sullivan and Karlseder, 2010). In metazoans, this highly conserved nucleoprotein structure is shortened during each cellular division cycle, and can also be damaged by oxidative stress, potentially accelerating the shortening process (Young 2018). Upon reaching a critical minimum length, replicative senescence is triggered in order to prevent mutation and cancer (Campisi and d’Adda di Fagagna 2007). For these reasons, telomere length is a good proxy for long-term organismal health and longevity (Shalev et al. 2013). Given that telomere length is influenced by environmental stress and predicts long-term condition, it is likely that telomere shortening may be involved in the generation of life-history trade-offs and carryover effects (Young 2018). To our knowledge, no study has investigated how UVBR exposure in early life might influence telomere lengths later in life. Improving understanding of how the dose, irradiance, and duration of UVBR exposure interact on such carryover effects is crucial for elucidating the impacts of complex and increasing natural UVBR exposures on long-term amphibian health (Hegglin and Shepherd 2009; Mckenzie et al. 2011; Olker et al. 2013).

Our aim was to investigate how exposure of larvae to different doses and irradiances of acute UVBR affects size, condition, performance, and relative telomere lengths post-metamorphosis. We acutely exposed tadpoles of the Australian green tree frog (*Litoria caerulea*, White 1790) to a combination of different UVBR irradiances and doses in a fully factorial laboratory experiment, and then reared these larvae to metamorphosis, assessing a suite of life-history and performance traits in the juvenile frogs upon development. UVBR irradiance determines the rate of DNA damage production (Londero et al. 2019), so we hypothesised that high irradiance UVBR would hinder successful metamorphosis, and would be more detrimental to growth, jumping performance and foraging efficiency in juvenile frogs than an equivalent dose of low irradiance UVBR. Given that animals exposed to high irradiance UVBR are expected to experience reduced growth and increased oxidative stress, both of which can shorten telomeres, we also hypothesised that this treatment would cause shorter relative telomere lengths in metamorphs.

## MATERIALS AND METHODS

### Ethics statement

This research was approved by The University of Queensland Animal Ethics Committee (approval no. SBS/089/19 and 2021/AE000365) and animal collection permission was granted by the Queensland Department of Environment and Science (permit no. WISP17421516).

### Study species

*Litoria caerulea* (White 1790) is a common species distributed throughout northern and eastern Australia. The IUCN lists this species as ‘Least Concern’, however population declines have been documented in some regions (Berger et al. 1998). *L. caerulea* is relatively resilient to UVBR (Lundsgaard et al. 2021), which is consistent with its ecology, laying transparent gel egg masses in unsheltered, ephemeral water bodies that are exposed to UVBR levels of up to 500 μW cm^−2^ (at water surface, van Uitregt et al. 2007).

### Animal collection and husbandry

Seven freshly laid *L. caerulea* egg masses were collected from flooded roadsides in southeast Queensland, Australia, and transported to The University of Queensland in plastic bags containing water from the collection site. Each clutch was maintained in a separate 5 L plastic tank of carbon-filtered Brisbane city tap water and housed at 22.5 ± 1°C on a 12L:12D photoperiod regime using non-UVBR fluorescent lights. At the commencement of UVBR exposure treatments four weeks post-laying, larvae (Gosner stage 25; Gosner 1960) were individually housed in clear plastic containers (1 L; 15 × 10 × 7 cm) filled to a depth of 5 cm with carbon-filtered Brisbane city water (see *Experimental Design*). Larvae remained in these containers following UVBR exposure and were reared to metamorphosis on thawed spinach, with half water changes (of carbon-filtered Brisbane city water) made twice per week to maintain water quality. When larvae developed front legs (Gosner stage 42), containers were partially emptied of water and angled such that metamorphosing animals could climb out of the water. Newly metamorphosed juvenile frogs were held individually in their original housing containers with perforated lids and soaked paper towel added as substrate. Frogs were fed crickets and cockroaches every four days until completion of tests one month post-metamorphosis, after which they were euthanised in a buffered Tricaine-S bath (MS-222; Aqua-Life, Nanaimo, Canada; 0.5 g L^−1^).

### Experimental Design

The experimental design employed in this study is also described in Lundsgaard et al. (2021) which reported on an earlier phase of this larger experiment. Four-week-old, individually-housed larvae (n = 160), were randomly allocated to one of ten UVBR treatments in a 3 × 3 factorial design (plus a no-UVBR treatment; n = 16 per treatment) that were centred around midday and administered in addition to the 12L:12D non-UVBR background lighting. Three levels of UVBR irradiance (low: 8.7 μW cm^−2^, medium: 35.5 μW cm^−2^, and high: 70 μW cm^−2^; Table S1) were fully crossed with three levels of UVBR dose (low: 1 day and approximately 2.5 kJ m^−2^ UVBR, medium: 4 days and approximately 10 kJ m^−2^ UVBR, and high: 8 days and approximately 20 kJ cm^−2^ UVBR; Table S1). To decouple dose and irradiance, the daily time interval of exposure was adjusted for each irradiance treatment so that daily dose remained constant (8, 4 and 1 hour of exposure per day for the low, medium and high irradiance treatments, respectively). Temperature during and after the UVBR exposure period was cycled daily to represent summer conditions at the site of collection, ranging from 21 – 31°C. Water temperatures of each treatment were measured during the experiment using waterproof iButton temperature data loggers and ranged from 21.5°C to 29°C. Additional temperature fluctuations associated with heat emitted by the UVBR lights were minimised by placing trays of ice on ‘buffer shelves’ immediately underneath the high irradiance UVBR treatment during exposure events.

After the acute UVBR exposures, larvae were reared to metamorphosis as described earlier (Fig. 1). Age (from egg), size (mass, snout-to-vent length, tibiofibular length and interorbital distance), and body condition (scaled mass index) were measured at metamorphosis (defined as the full reabsorption of the tail at Gosner stage 46). Growth rate of juvenile frogs in the first month post-metamorphosis was also determined. One month after metamorphosis, juvenile frogs were tested for jumping performance and foraging efficiency. Upon completion of tests, frogs were euthanised in a MS-222 bath and carcasses were immediately stored at −80°C for analysis of relative telomere lengths (see *Traits* section for more details).

**Figure 1.**
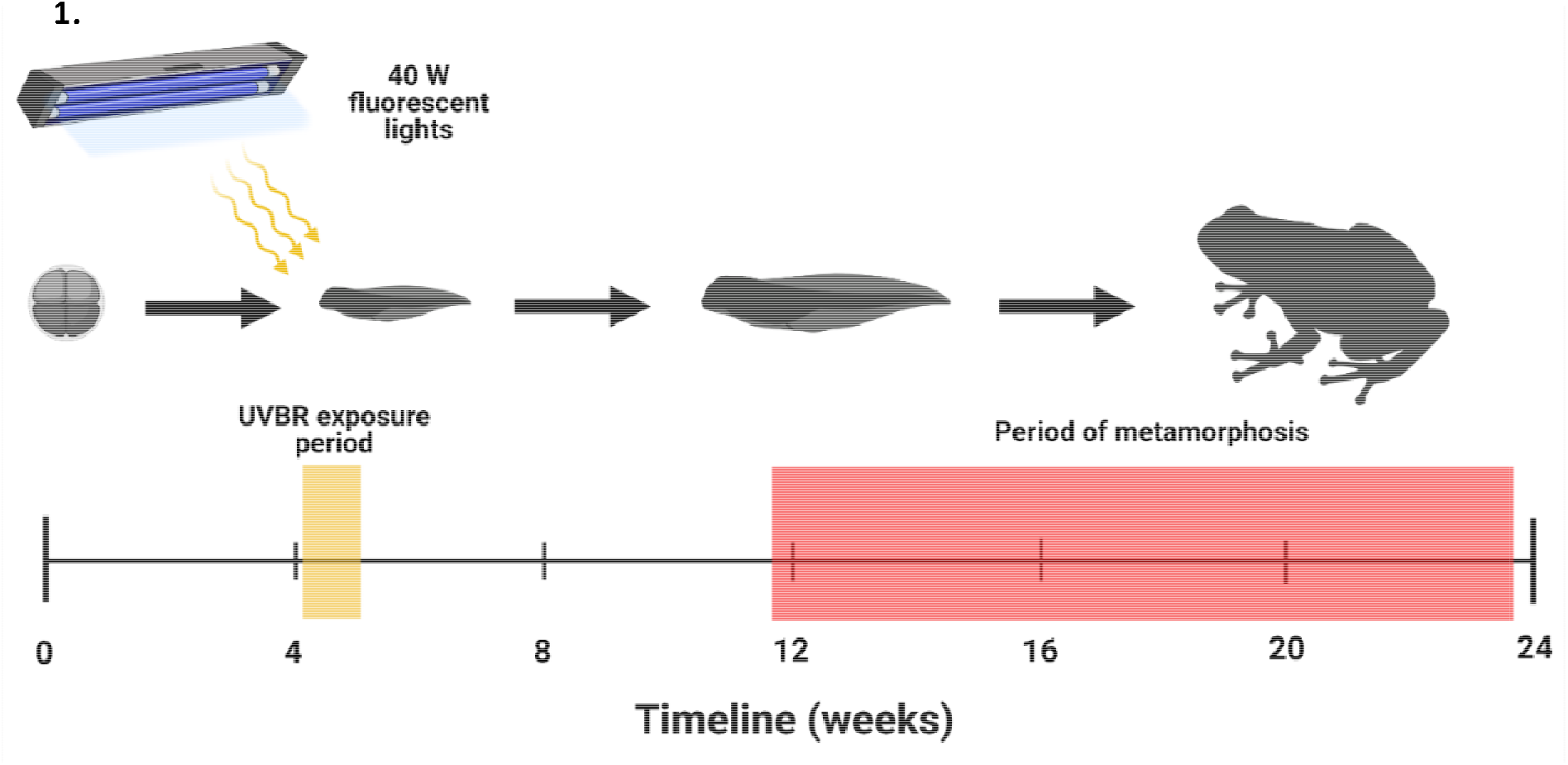
A schematic of the experimental timeline (in weeks). The yellow bar on the timeline represents the period of exposure of *L. caerulea* larvae (Gosner stage 25) to UVBR treatments, whilst the time period in which metamorphosis occurred is demonstrated by the red bar. The amphibian silhouettes are for illustrative purposes only. Created with BioRender.com.

#### UVBR Exposures

UVBR conditions were generated using 1.2 m, 40 W fluorescent light tubes (no UVBR = 0 zero bulbs; low irradiance = two bulbs at 51 cm; medium irradiance = six bulbs at 27 cm; high irradiance = eight bulbs at 12 cm; Repti-Glo 10.0, Exo Terra, Montreal, Canada). UVBR and ultraviolet-A (UVAR; 315 – 400 nm) levels at the water surface of each container were measured using a radiometer/photometer (IL1400BL, International Light Inc., Newburyport, USA) to ensure consistent levels across experimental shelves.

Fluorescent lights are a suitable substitute for natural sunlight when housing reptiles and amphibians, emitting important biologically active wavelengths across the ultraviolet (UV), visible and infrared spectra (Baines et al. 2016). That said, the physiological effects of artificial lighting cannot be directly extrapolated to sunlight exposure because of differences in spectral composition (Baines et al. 2016). Daily, natural fluctuations of UVBR and UVAR irradiances are correlated (Schuch et al. 2015b), so the ratio of UVBR:UVAR was kept as similar as possible between treatments. UVBR levels used in this study are much lower than ambient UVBR levels measured in the region (500 μW cm^−2^ in air at water level at midday in Brisbane, Australia; van Uitregt et al. 2007), to help account for attenuation by cloud cover, vegetation cover and dissolved organic matter that reduce aquatic UVBR levels in the wild (Palen et al. 2002; Palen and Schindler 2010; Olker et al. 2013; Alton and Franklin 2017).

## Traits

### Age, size and condition at metamorphosis

Developing larvae were monitored daily, with age at metamorphosis defined as the number of days between egg laying and full reabsorption of the tail (Gosner 46). Juvenile frogs were weighed and photographed on the day of metamorphosis and again one month later, allowing for calculations of post-metamorphic growth. Snout-to-vent length, leg length (tibiofibular bone) and interorbital distance (length between the eyes) were determined from photographs (iPod touch 5^th^ generation, Apple, California, USA) of the dorsal surface of each frog (with a ruler for scale) using the software program *ImageJ* (National Institutes of Health, Bethesda, Maryland, USA).

Of the three linear size metrics measured, interorbital distance correlated most strongly with body mass and was therefore used in mass/length calculations of body condition, as suggested by Peig and Green (2009). Condition factor of juvenile frogs was calculated using the scaled mass index (SMI) method developed by Peig and Green (2009), which has been demonstrated as the most suitable and accurate condition index across a range of taxa (Peig and Green 2009, 2010; Brodeur et al. 2020). In accordance with Brodeur et al. (2020) who confirmed the applicability of this method for use in amphibians, we defined the scaling exponent *b* through a non-linear power function regression (mass = *α*[interorbital distance]^*b*^) for the sample population (minus two outliers) which was: *γ* = 0.0012*x*^2.9624^ (R^2^ = 0.793), to obtain size-independent SMI values. This scaling exponent conforms with allometric scaling observed in other taxa, which typically ranges between 2.5 and 3.2 (Green 2001). Two outliers did not fit the mass/length trend of the sample population (n = 50) and were therefore removed from calculation of the regression, following Peig and Green (2009). SMI was calculated relative to the average interorbital distance of the sample population (7.32 mm), giving the estimated mass that each frog (including the two outliers) would have at this fixed body size. Larger SMI values thus indicate larger energy reserves and provide an effective estimate of body condition (Peig and Green 2009).

#### Foraging efficiency

Four weeks post-metamorphosis, and following a four-day fasting period, juvenile frogs were tested for indices of foraging efficiency. Individual frogs were placed under a clear lid in the middle of a rectangular foraging arena (17 × 30 × 10 cm) for a five-minute adjustment period prior to testing. A cricket of known mass (7 – 34 mg; average cricket to frog mass ratio = 1:27, range = 1:104 – 1:10) was then placed under a holding container at the opposite end of the arena so that the frog was directly facing the cricket. Both holding containers were then lifted simultaneously and the frog was allowed to freely hunt the cricket in the hunting arena. The hunt was timed and recorded using a GoPro Hero 5 (at 120 fps and 1080p) and was terminated the moment the frog captured and swallowed the prey, or in some instances ‘gave up’ on the hunt due to fatigue (defined as the point in time in which the frog stopped eliciting stalking behaviour for 30 s or more). Frogs were then returned to the middle of the arena, placed under a lid, and rested for five minutes. The procedure was repeated two more times (three hunts per day), and these foraging efficiency tests were repeated again four days later. Videos were played back (Tracker Video Analysis and Modelling Tool, Open Source Physics) frame-by-frame by digitising the snout to determine the maximum successful strike distance and strike speed (leading to prey capture), as well as the time in pursuit of prey until capture. Only the longest and fastest strike was analysed, while the number of strikes and time in pursuit data were averaged across the six trials per animal. Individual trials where no stalking/hunting behaviour was elicited were excluded from analyses. Two animals did not elicit any stalking/hunting behaviour during the trials, instead attempting to escape the foraging arena, or not responding to the presence of prey. Thus, these animals were excluded from analysis.

#### Jumping performance

Four weeks post-metamorphosis, and two days after the first foraging efficiency test, the same juvenile frogs were tested for jumping performance. Frogs were placed on top of laminated grid paper and filmed immediately (GoPro Hero 5, 240 fps at 720p) while maximum escape behaviour was being elicited. Frogs that remained stationary were urged to jump by gentle contact to the vent. After at least five jumps, frogs were returned to the middle of the grid paper, placed under a lid, and rested for five minutes. The jump tests were then repeated once more. Recordings were played back (Tracker Video Analysis and Modelling Tool, Open Source Physics) frame-by-frame by digitising the snout to determine the maximum jump length and speed. The longest and fastest jump recorded was considered as the maximum jumping performance of the animal and used for analysis. If frogs jumped further or faster in a foraging efficiency trial, then these jumps were considered as the maximum jumping performance of the animal and used in the jumping performance analysis. Two animals did not elicit measurable jumps despite repeated attempts and were excluded from analysis.

#### Relative telomere lengths by qPCR

The qPCR-based relative telomere length assay followed Burraco et al. (2017). Approximately 10 mg (6-17 mg) of hind-leg muscle tissue was dissected (on ice) from each juvenile frog carcass (n = 2 – 8 per treatment). Genomic DNA was extracted from the finely chopped muscle tissue using a PureLink Genomic DNA Mini Kit (Invitrogen, 5791 Van Allen Way Carlsbad, CA, USA), in accordance with the manufacturer’s protocol. DNA concentrations were then determined using a Qubit fluorometer (Invitrogen, CA, USA, Cat # Q32857) and a Qubit dsDNA Broad-Range Assay Kit (Invitrogen, CA, USA, Cat # Q32853), in accordance with the manufacturer’s protocol. DNA samples were aliquoted and stored at −80°C until use.

The broadly conserved nature of telomeric sequence repeats in vertebrates allowed us to use the same primer set as Burraco et al. (2017), i.e. F: 5-AACCAGCCAAGTACGATGACAT-3′ and R: 5′-CCATCAGCAGCAGCCTTCA-3. Species-specific Quantitative PCR (qPCR) primers against the glyceraldehyde-3-phosphate dehydrogenase (GAPDH) housekeeping gene were designed using PrimerQuest (Integrated DNA Technologies, IA, USA) with acceptance of the default parameters, except that amplicon length was set to 95-105 bp. The GAPDH primers were: F: 5′CGGTTTGTTTGGGTTTGGGTTTGGGTTTGGGTTTGGGTT-3′, and R: 5′-GGCTTGCCTTACCCTTACCCTTACCCTTACCCTTACCCT-3′. qPCR was performed using iTaq Universal SYBR Green Supermix (Bio-Rad Laboratories Inc.). We used a 1:40 dilution of DNA samples to suit both the telomere and GAPDH primer amplification rates. qPCR for telomere and GAPDH genes were performed on separate plates, but in corresponding wells for each sample. We used 1.75 ng of genomic DNA per reaction, and primer concentrations of 500 nM in a 20 μL reaction containing 10 μL 2× SYBR Green Supermix. PCR cycles for amplification of telomeric repeats were 5 min at 95°C, and then 30 cycles of 1 min at 56°C and 30 s at 95°C. For GAPDH, the 5 min incubation at 95°C was followed by 40 cycles of 1 min at 60°C and 95°C for 30 s. All qPCR cycles were conducted on a Bio-Rad CFX Connect Real-Time System. The efficiency of each qPCR plate performed was determined from a standard curve by serially diluting a combined pool of all the samples (4-fold dilutions to 1:1024 concentration, and corresponding to 70.6, 17.65, 4.413, 1.103 and 0.276 ng of DNA per well). Samples were run in duplicate, and each assay included a separate standard curve as well as a no-template control (in triplicate) at the start and end of the plate (to detect any contamination associated with pipetting).

All PCR efficiencies were above 95%, and unique dissociation curves were produced in all assays. Contamination in no-template control wells (Ct value of 30, relative to telomere Ct values of approximately 16.3) could not be avoided despite all efforts. Vasilishina et al. (2019) notes that contamination is not uncommon for this method, and that this should not affect results of relative telomere length evaluation. Data were collected using Bio-Rad CXF Manager software (version 3.1, Bio-Rad), and results were exported to Excel. Relative telomere length was quantified from averaged Ct values in accordance with Vasilishina et al. (2019) by normalising to the reference DNA sample: ΔΔ*Ct* = Δ*Ct_sample_* − Δ*Ct_reference_*, where Δ*Ct* = *Ct_TEL_* − *Ct_GAPDH_* for each sample. ΔΔ*Ct* values were used in statistical analysis and fold change in telomere length relative to the reference DNA sample was calculated for graphical display, where fold change = 2^−ΔΔ*Ct*^ (Pfaffl 2001).

#### Statistical Analysis

Statistical analyses were performed using R version 3.4.2 (“*Short Summer*”; R Core Team 2017). Data are presented as means ± standard error, unless otherwise stated, and egg clutch was included as a random effect in all models where it accounted for significant variance. Where interactive effects were non-significant, they were systematically removed to generate the simplest model representative of the data. The no-UVBR treatments proved detrimental to larval fitness for this species (Lundsgaard et al. 2021), resulting in insufficient metamorphs for statistical analysis (Fig. 1). Therefore, the no-UVBR treatment was excluded from analyses.

Treatment-specific effects on the proportion of animals metamorphosing were modelled with a binomial generalised linear model (GLM; function *glm*), whilst average age at metamorphosis was modelled with a linear mixed effects (LME) model (*lmerTest* package, function *lmer*; Kuznetsova et al. 2017). Additionally, a holistic analysis of treatment effects on the progression of metamorphosis over time was assessed with a Cox mixed effects survival analysis utilising the *coxme* (Therneau, 2018), *Matrix* (Bates and Maechler 2017) and *survival* (Therneau and Grambsch 2000; Therneau 2015) packages.

The correlation between age and mass at metamorphosis was assessed with a simple linear regression of the whole sample population. Treatment-specific differences in body mass and SMI of metamorphs were modelled with LME models. Following Alton and Franklin (2012), the correlation matrix of body mass, snout-to-vent length, interorbital distance, and leg length was assessed with a principal components analysis (PCA) to investigate treatment-associated differences in overall body size at metamorphosis. Of the resulting morphological variables generated, principal component (PC) 1 accounted for 79.43% of the variation, and PC2 accounted for 11.17% of the variation. The PC factor scores for PC1 were then modelled with an LME model. PC2 did not explain sufficient variation to be of particular interest (Quinn and Keough 2002), and so was not assessed. To assess treatment-associated differences in the growth of juvenile frogs in terms of overall body size, a PCA was performed using the same morphological measurements at one month post-metamorphosis. PC1 accounted for 86.28% of the variation and was used in analysis, while PC2 did not explain sufficient variation to be retained for further analysis (7.02%). The PC1 factor scores were modelled with an LME model with the PC1 factor scores from the initial size-at-metamorphosis data included as a covariate.

The data for prey capture time as well as average and maximum strike attempts needed for successful prey capture were skewed, and therefore modelled with negative binomial GLMs (*Mass* package, function *glm.nb*; Venables and Ripley 2002) with frog mass included as a covariate. Cricket mass was also included as a covariate in the analysis of maximum strike attempts until prey capture. Maximum distance and speed for both jumping performance and successful prey strikes were analysed with LME models. Frog leg length (tibiofibular length) was included as a covariate for jumping performance models, while frog mass was a better predictor in models of maximum strike distance and speed. ΔΔ*Ct* values (a proxy for telomere lengths) were also modelled with an LME model with frog mass as a covariate. Tukey contrasts were conducted post hoc using the *multcomp* package (function *glht*; Hothorn et al. 2008).

## RESULTS

The survival analysis revealed a significant effect of UVBR dose on the progression of metamorphosis of *L. caerulea* larvae over time (χ^2^_[2]_ = 7.742, *P* = 0.021), with less than half as many juvenile frogs developing in the low dose treatments compared to the medium and high dose treatments (*Z* = 2.170, *P* = 0.076; *Z* = 2.602, *P* = 0.025, respectively; Fig. 2A). However, there was no significant difference in the average age at metamorphosis of these individuals (*F_2,42_* = 0.099, *P* = 0.906). There was no interaction between UVBR dose and irradiance on the progression or average age at metamorphosis (χ^2^_[4]_ = 3.785, *P* = 0.436; *F_4,36_* = 2.012; *P* = 0.113, respectively), and no significant main effect of UVBR irradiance on these metrics (progression of metamorphosis: χ^2^ = 4.970, *P* = 0.083; age at metamorphosis: *F_2,41_* = 0.906, *P* = 0.412, respectively; Fig. 2B).

**Figure 2.**
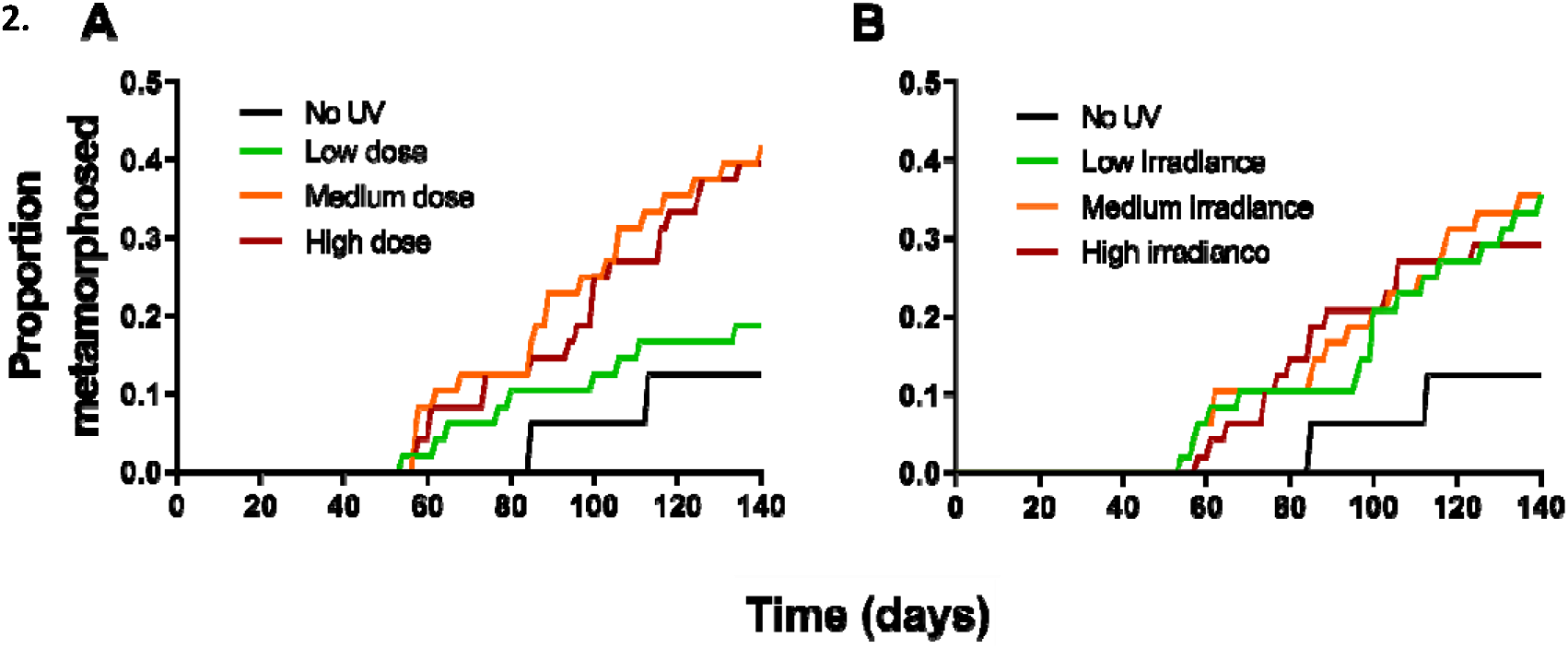
Effect of UVBR dose (A) and irradiance (B), on the progression of metamorphosis (as a proportion) of *L. caerulea* larvae over a 140-day period following commencement of treatments. No interaction between dose and irradiance was present, so data for the dose treatments are pooled across irradiances, and vice versa, to observe main effects (n = 48 per treatment level). The no-UVBR control treatment (black line) is provided in both panels for reference (n = 16).

There was a moderate positive correlation between age and mass at metamorphosis (*F_1,48_* = 16.54, *P* < 0.001; R^2^ = 0.256, Fig. 3), with age at metamorphosis a strong predictor of metamorph mass in the LME model (*F_1,40_* = 17.641, *P* < 0.001). There was no interactive effect of UVBR dose and irradiance on mass or overall size at metamorphosis (*F_4,34_* = 0.249, *P* = 0.908; *F_4,36_* = 0.220, *P* = 0.926, respectively), and no main effect of UVBR dose on these metrics (mass: *F_2,39_* = 1.636, *P* = 0.208; overall size: *F_2,41_* = 2.543, *P* = 0.091; Fig. 4A). However, there was a marginal effect of UVBR irradiance on mass at metamorphosis (*F_2,39_* = 2.925, *P* = 0.066). Given that UVBR irradiance and dose did not significantly affect age at metamorphosis (see previous paragraph), we justified the removal of this covariate from analysis to prevent confounding effects with mass, which in turn yielded a significant effect of UVBR irradiance, but still not dose, on mass at metamorphosis (irradiance: *F_2,38_* = 4.094, *P* = 0.025; dose: *F_2,39_* = 1.761, *P* = 0.185, respectively) and overall size at metamorphosis (irradiance: *F_2,43_* = 3.504, *P* = 0.039; dose: *F_2,43_* = 2.772, *P* = 0.074). Larvae exposed to high irradiance UVBR metamorphosed with 20% less mass, on average, than larvae exposed to the low and medium irradiance treatments (*Z* = −2.617, *P* = 0.024; *Z* = −2.443, *P* = 0.039, respectively; Fig. 4B). Size at metamorphosis in turn predicted body size in the month following (*F_1,38_* = 96.867, *P* < 0.001), but there was no significant effect of larval UVBR treatment on growth during this time (dose: *F_2,37_* = 0.305, *P* = 0.739, Fig. 4C; irradiance: *F_2,37_* = 0.784, *P* = 0.464, Fig. 4D; interaction: *F_4,36_* = 1.203, *P* = 0.326).

**Figure 3.**
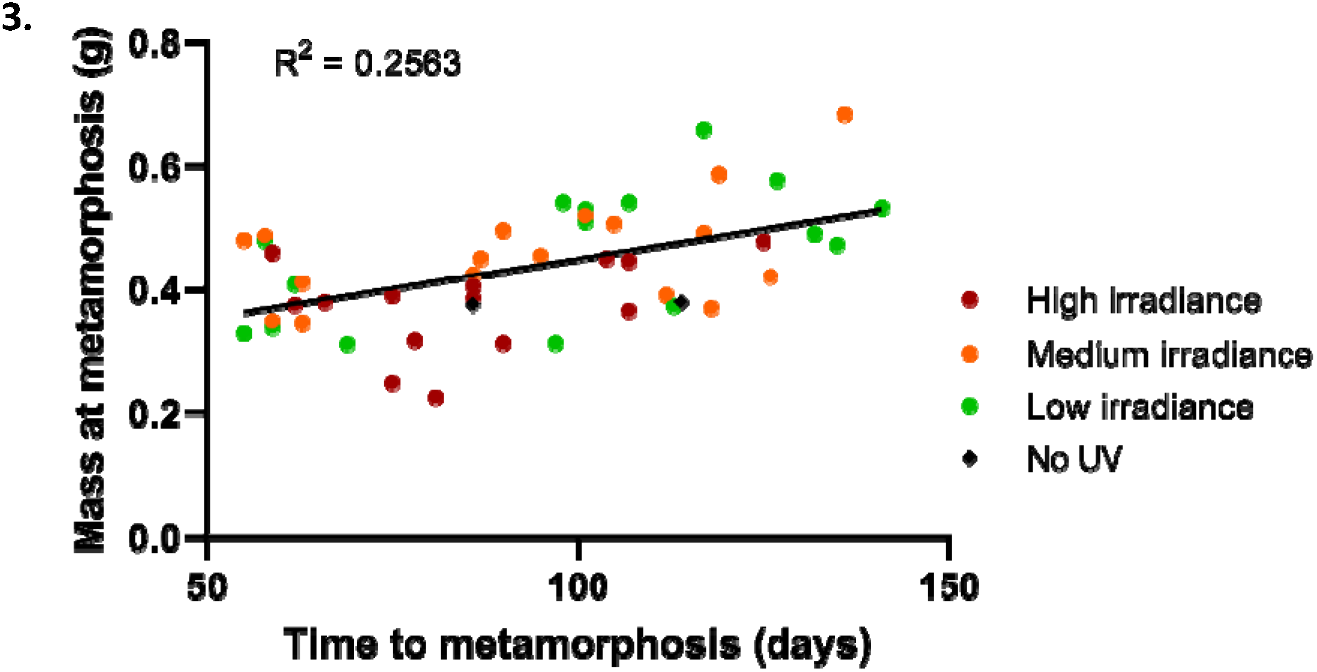
Correlation between age at metamorphosis and mass at metamorphosis (g) of *L. caerulea* larvae. Data points represent individual animals, color-coded by UVBR irradiance treatment (n = 14 – 17 per irradiance, pooled across UVBR doses), with no-UVBR control animals indicated by black points (n = 2). Trend line indicates a moderate positive correlation between time to metamorphosis and mass (R^2^ = 0.2563).

**Figure 4.**
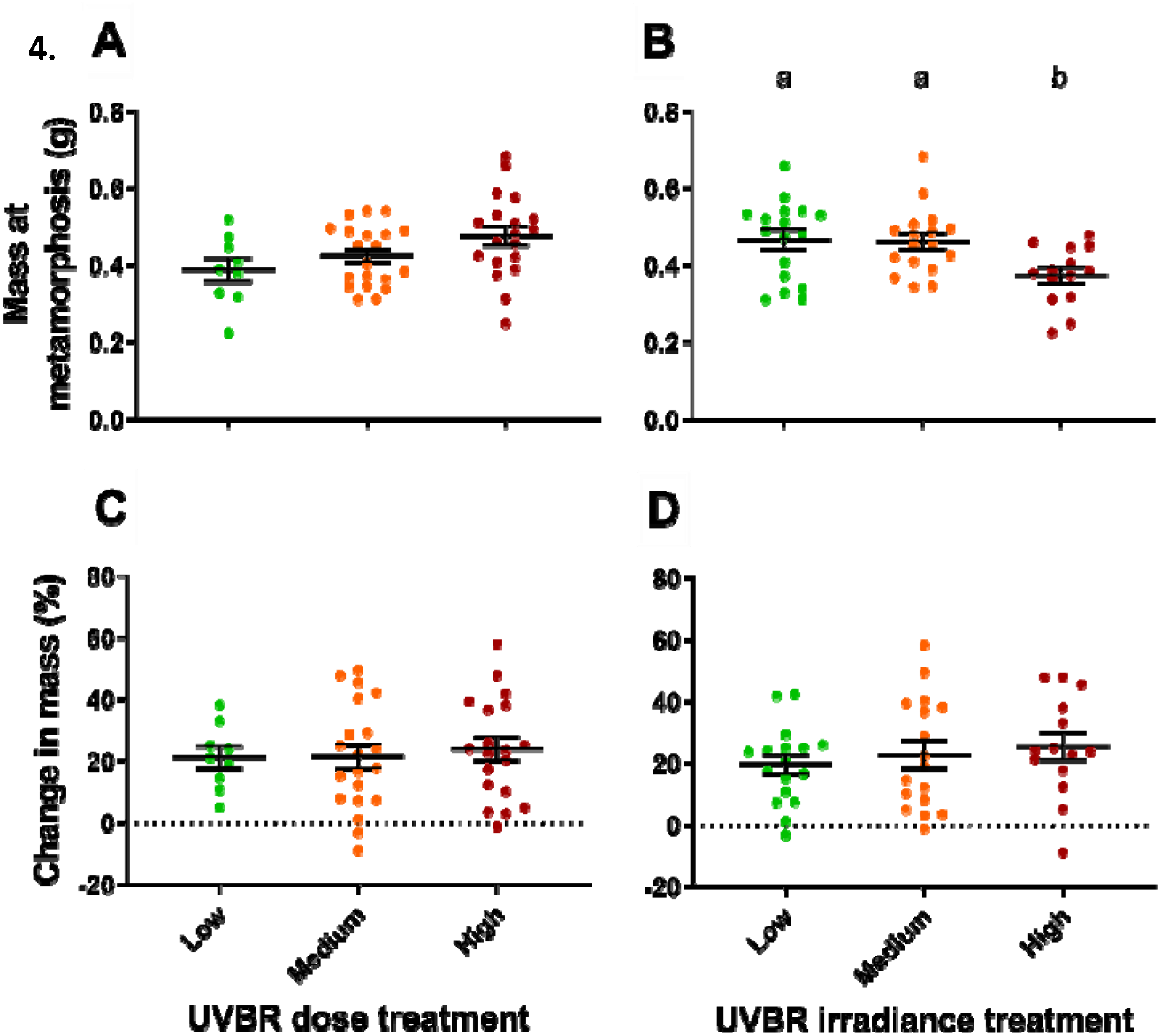
Effect of UVBR dose and irradiance (pooled) on mass at metamorphosis (g; A and B, respectively) and growth after metamorphosis (C and D, respectively) of *L. caerulea* metamorphs (n = 9-20 per pooled treatment level). Growth after metamorphosis was measured as a percentage change in mass after 30 days, with no net change in mass indicated by the grey dotted line. Data are presented as means ± s.e.m., and lowercase letters denote significant differences (*P* < 0.05) between treatment groups.

There was a significant interaction between UVBR dose and irradiance on juvenile frog body condition (interaction: *F_4,36_* = 2.797, *P* = 0.040), whereby the effect of UVBR dose depended on the irradiance at which it was administered. High UVBR dose exposure improved metamorph body condition when administered at a low irradiance but reduced body condition when administered at medium irradiance (Fig. 5). Larvae exposed to high irradiance UVBR had reduced body condition at metamorphosis regardless of the dose administered, with an average 7.6% reduction in mass when scaled for the average interorbital distance of the sample population, compared to the low and medium irradiance treatments. Because of these interactive effects, the low dose UVBR treatment yielded the greatest differences in body condition depending on the irradiance of administration, with larvae that received a low UVBR dose metamorphosing with the worst SMI when administered at a low irradiance, but the best SMI when administered at a medium irradiance (*t*_35_ = −2.856, *P* = 0.019; Fig. 5).

**Figure 5.**
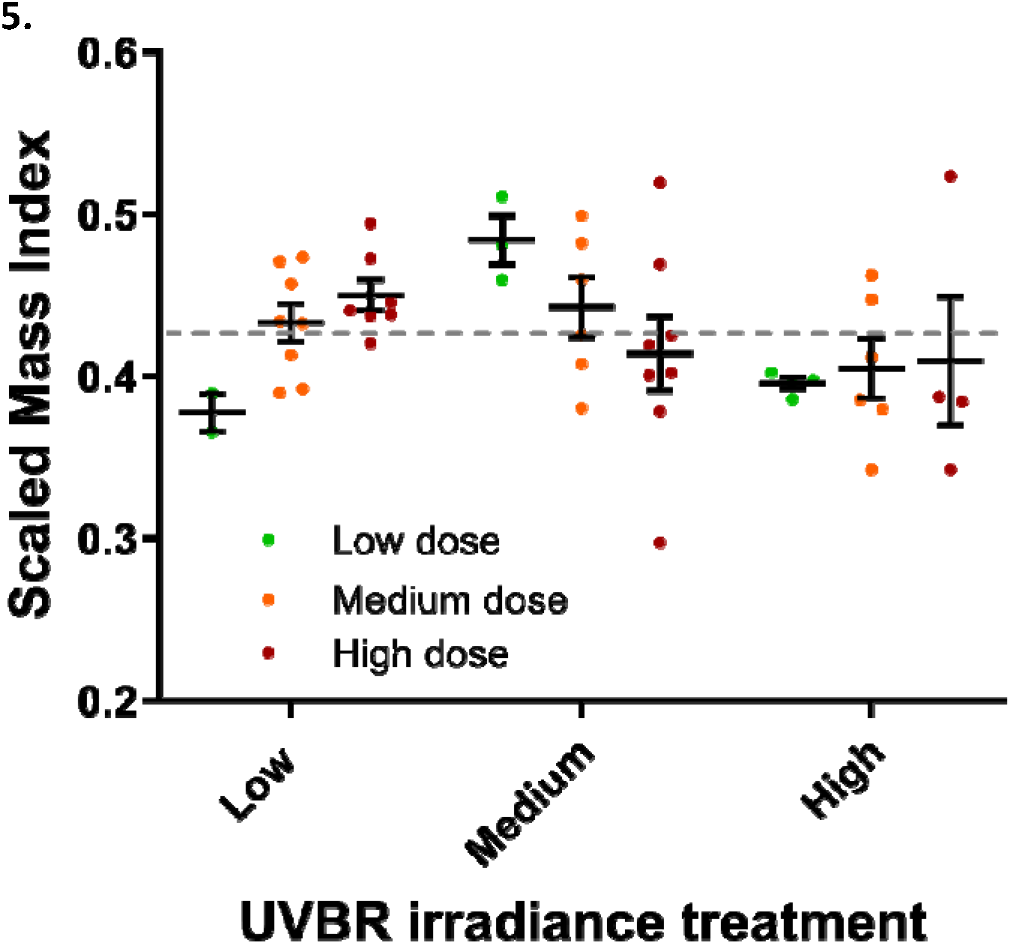
Effect of UVBR dose (colours) and irradiance (columns) on body condition (scaled mass index method; SMI) of *L. caerulea* metamorphs (n = 2-8 per treatment). SMI values represent the estimated mass (g) when scaled to the mean interorbital distance of the sample population. Data points represent individual animals, and error bars indicate treatment means ± s.e.m. The grey dashed line represents the mean SMI of the sample population, such that points above the line indicate good relative body condition, while points below the line indicate poor relative body condition.

There was no significant effect of larval UVBR treatment on foraging efficiency of subsequent juvenile frogs, as measured by foraging time (dose: χ^2^_[2]_ = 0.408, *P* = 0.816; irradiance: χ^2^_[2]_ = 0.383, *P* = 0.826; interaction: χ^2^_[4]_ = 2.241, *P* = 0.692), average strike attempts (dose: χ^2^_[2]_ = 1.573, *P* = 0.455; irradiance: χ^2^ = 0.449, *P* = 0.799; interaction: χ^2^ = 0.450, *P* = 0.978), maximum strike attempts (dose: χ^2^ = 1.718, *P* = 0.424; irradiance: χ^2^ = 0.215, *P* = 0.898; interaction: χ^2^_[4]_ = 1.098, *P* = 0.895), maximum successful strike distance (dose: *F_2,37_* = 0.995, *P* = 0.379; irradiance: *F_2,37_* = 1.151, *P* = 0.327; interaction: *F_4,37_* = 1.795, *P* = 0.151) and maximum successful strike speed (dose: *F_2,37_* = 0.409, *P* = 0.667; irradiance: *F_2,37_* = 1.274, *P* = 0.292; interaction: *F_4,37_* = 1.066, *P* = 0.387; Table 1). Cricket size did not influence maximum strike attempts until successful prey capture (χ^2^ = 0.081, *P* = 0.775). However, there was a significant effect of frog mass on foraging efficiency as measured by maximum successful strike distance and strike speed (*F_1,37_* = 13.338, *P* < 0.001; *F_1,37_* = 9.435, *P* = 0.004, respectively), with larger frogs able to jump farther and faster when striking prey (Fig. 6).

**Table 1.**
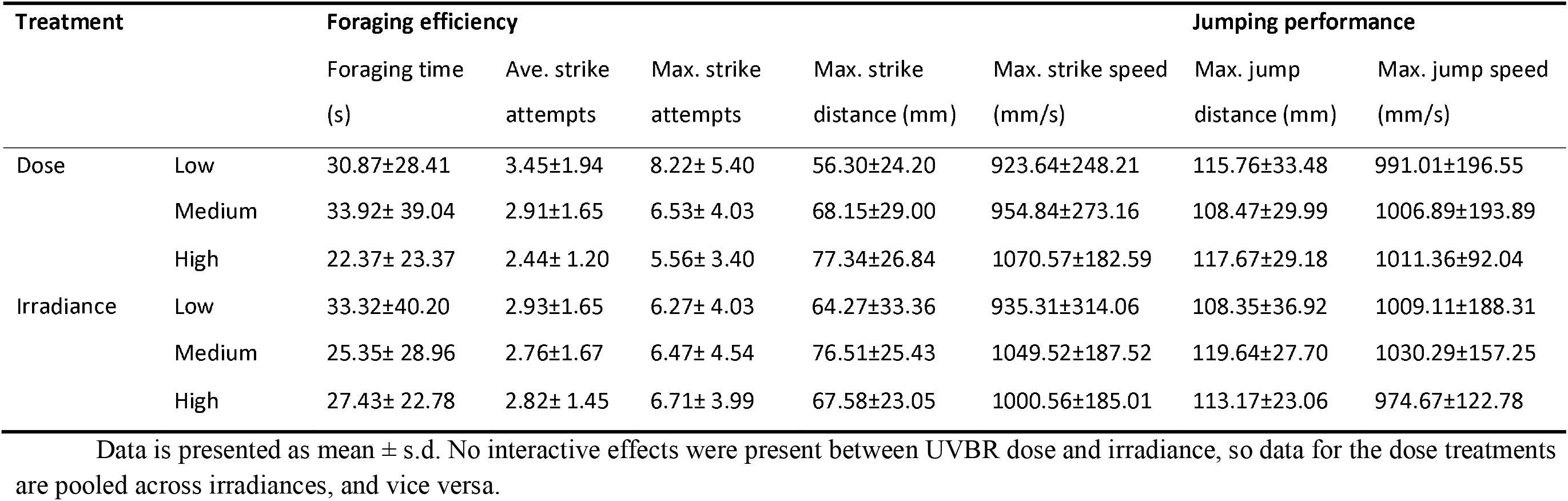
Performance data for *L. caerulea* metamorphs in each UVBR treatment. No statistically significant treatment effects occurred. Data is presented as mean ± s.d. No interactive effects were present between UVBR dose and irradiance, so data for the dose treatments pooled across irradiances, and vice versa.

**Figure 6.**
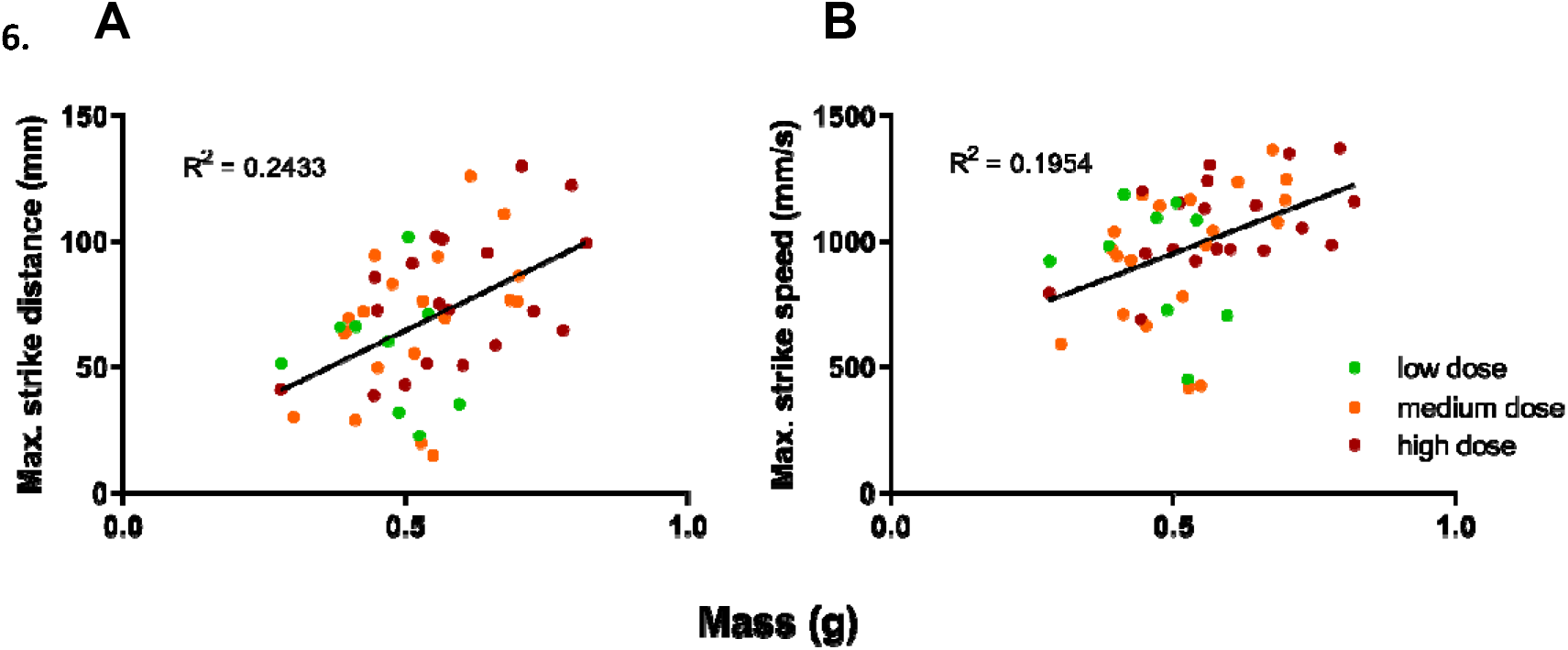
Correlation between mass (g) of *L. caerulea* frogs 1-month post-metamorphosis, and foraging performance metrics (maximum strike distance [mm] (A) and speed [mm/s] (B) of successful prey capture). Data points represent individual animals, color-coded by UVBR dose treatment (n = 9 – 20 per irradiance, pooled across UVBR doses). Trend lines indicates a moderate positive correlation between mass and foraging performance.

There was no significant effect of larval UVBR treatment on metamorph jumping distance (dose: *F_2,33_* = 0.130, *P* = 0.879; irradiance: *F_2,32_* = 2.099, *P* = 0.139; interaction: *F_4,33_* = 1.113, *P* = 0.367) and maximum jumping speed (dose: *F_2,38_* = 0.377, *P* = 0.689; irradiance: *F_2,38_* = 0.250, *P* = 0.780; interaction: *F_4,38_* = 0.202, *P* = 0.936; Table 1). However, there was a significant effect of frog size (especially tibiofibular length) on jumping performance as measured by maximum jumping distance and maximum jump speed (jumping distance: *F_1,37_* = 7.789, *P* = 0.008; jumping speed: *F_1,38_* = 13.442, *P* < 0.001), with larger frogs able to jump farther and faster (data not shown)

UVBR irradiance had a significant effect on relative telomere lengths (*F_2,38_* = 5.573, *P* = 0.008; Fig. 7B), with juvenile frogs exposed to medium irradiance UVBR as larvae having shorter relative telomere lengths as frogs than those previously exposed to low and high irradiance UVBR treatments (*Z* = 3.116, *P* = 0.005; *Z* = −2.524, *P* = 0.031, respectively). There was no interaction between larval UVBR dose and irradiance on relative telomere lengths (*F_4,33_* = 0.524, *P* = 0.719), and no main effect of UVBR dose on this metric (*F_2,38_* = 0.146, *P* = 0.865; Fig. 7A). Frog mass (at the time of sampling) did not have a significant effect on relative telomere lengths (*F_1,38_* = 0.874, *P* = 0.356).

**Figure 7.**
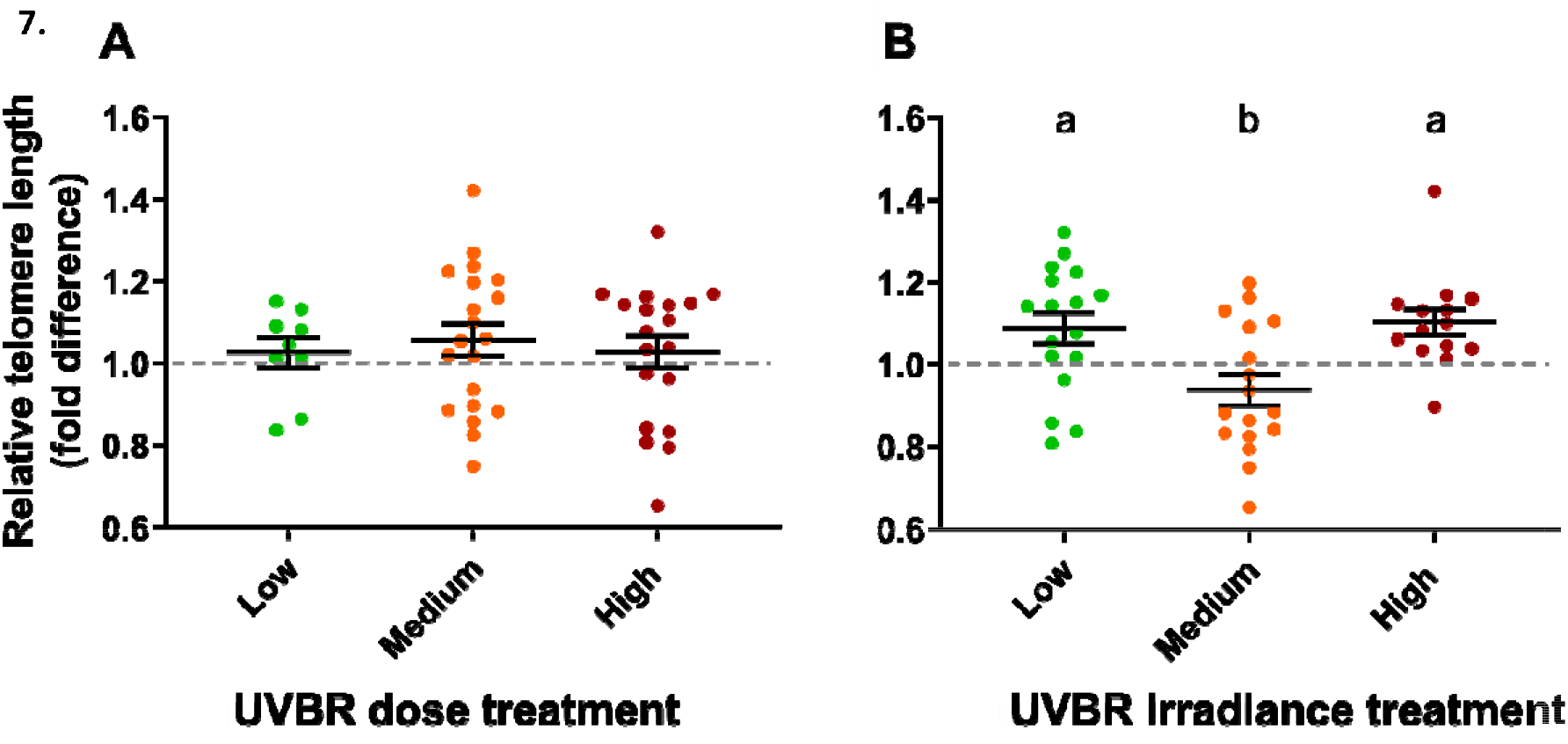
Effect of UVBR dose (A) and irradiance (B) pooled across the levels of the other factor, on relative telomere lengths of *L. caerulea* juveniles 1 month post-metamorphosis (n = 9-20 per pooled treatment level), as a fold difference from the reference sample (a pool of all samples, indicated by the grey dashed line). Data are presented as means ± s.e.m., and lowercase letters denote significant differences (*P* < 0.05) between treatment groups.

## DISCUSSION

We present three major findings in this study. We found that dose and irradiance of acute UVBR exposure in larvae both influence long-term carryover effects into metamorphosis in a trait-specific manner. High irradiance UVBR had contrasting effects across different life-history stages (Lundsgaard et al. 2021), with evidence for life-history trade-offs negatively impacting condition at metamorphosis. Finally, we demonstrated empirical evidence of telomere shortening associated with UVBR exposure. These findings contribute to our understanding of how acute UVBR exposure regimes in early life affect later life-history stages in amphibians, further elucidating the role of this pervasive stressor in shaping amphibian population dynamics.

In this study, four-week-old *L. caerulea* larvae (Gosner stage 25) exposed acutely to UVBR experienced long-term carryover effects into the juvenile stage, driving changes in metamorphosis, body condition and relative telomere lengths. Only a few other studies have explicitly demonstrated carryover effects of UVBR exposure in amphibians through metamorphosis (Pahkala et al. 2001, 2003; Ceccato et al. 2016). Pahkala et al. (2001) found that embryos exposed to an enhanced dose of UVBR suffered from a higher prevalence of developmental abnormalities as larvae, were delayed in metamorphosis, and were smaller as juveniles compared to animals that were not exposed to UVBR. Ceccato et al. (2016) found that a six-week exposure to UVBR in the larval phase led to immune system changes in subsequent juvenile frogs. Our work corroborates the pervasive nature of UVBR-induced carryover effects in amphibians through metamorphosis, highlighting the need to consider such long-term effects when modelling UVBR effects on amphibian populations.

The medium and high UVBR dose treatments significantly improved progression into metamorphosis compared to the low dose treatment, with reduced mortality throughout the larval phase and during the transition through metamorphosis. These results support the evidence for beneficial effects of high UVBR doses in this species, which include improved growth, development, and swimming performance of larvae (Lundsgaard et al. 2021). These results may reflect increased endogenous vitamin D_3_ synthesis in larvae and frogs previously exposed to high UVBR doses, a vitamin essential for maintaining calcium homeostasis, which in turn affects bone mineralisation, muscular function and nerve function (Stiffler 1993; Antwis and Browne 2009; Michaels et al. 2015). It is possible that a deficiency in calcium stores hindered successful metamorphosis in animals that did not receive sufficient UVBR exposure, because of the need for increased bone formation during this time (Stiffler 1993).

Although the provision of high irradiance UVBR has been shown to benefit larval swimming performance in this species (Lundsgaard et al. 2021), these benefits of high irradiance UVBR did not carryover to foraging and jumping performance post-metamorphosis. In fact, larval exposure to high irradiance UVBR had mainly detrimental carryover effects, resulting in reduced size and poorer body condition at metamorphosis, both of which are associated with reduced fitness and survival in amphibians (Chelgren et al. 2006; Scott et al. 2007). The positive correlation between mass and some metrics of foraging efficiency in this study lend further support to the detrimental fitness effects of reduced size at metamorphosis. As with previous studies of UVBR-induced carryover effects on amphibian metamorphosis, the detrimental effects of high UVBR irradiance were not evident in the life-history stage being exposed (Lundsgaard et al. 2021), but instead manifested post-metamorphosis (Pahkala et al. 2001; Ceccato et al. 2016). These results highlight the importance of tracking fitness consequences of UVBR exposure into later life-history stages, as such latent effects may otherwise be missed. The contrasting effects of larval exposure to high irradiance UVBR across different life stages is evidence for a life-history trade-off that may be driven by changes in energy balance (Pechenik 2006). It is possible that the increased energy requirements for somatic maintenance in animals exposed to high irradiance UVBR as larvae may have led to reduced energy reserves being available at metamorphosis, as indicated by the lower SMI values (Alton et al. 2012). Such UVBR-induced life-history trade-offs have been observed in other taxa (Debecker et al. 2015).

The mechanisms driving the somewhat opposing effects of UVBR dose and irradiance on juvenile *L. caerulea* health are not known. However, it seems that certain post-metamorphic benefits conferred by a high dose of UVBR in the larval phase were only expressed at lower, more tolerable irradiances. This was particularly apparent with the SMI data. Although caution is warranted in interpretation given the small sample size, our data show a positive trend between metamorph body condition and UVBR dose when administered at a low irradiance. At medium irradiance, beneficial effects of UVBR exposure are only conferred at a low and medium dose, whilst any amount of exposure to high irradiance UVBR proved detrimental to metamorph body condition. These results suggest that the effect of a given UVBR dose are highly dependent on the irradiance at which it is administered at, which highlights the importance of the rate of DNA damage production (determined by irradiance) for physiological outcomes (Pandelova et al. 2006; Londero et al. 2019). It seems that the detrimental effects of even large doses can be managed if irradiance, and the rate of formation of pyrimidine dimers in DNA, is low. If however, UVBR irradiance is great enough that the rate of DNA damage exceeds the rate of DNA repair (dose-toxicity threshold), then an accumulation of DNA damage is expected (Pandelova et al. 2006). This accumulation of damage, or the rate at which it is induced, along with the energy required to repair it, could explain the detrimental effects of the high irradiance UVBR treatments in later life-history stages (Alton et al. 2012; Debecker et al. 2015). Namely, this molecular effect is likely to influence energy use, oxidative stress, and even epigenetic modifications and telomere lengths in exposed animals, all of which have been implicated as generators of carryover effects (O’Connor et al. 2014; Young 2018).

Although high irradiance UVBR had the strongest impact on short-term fitness indices of *L. caerulea* at metamorphosis, it was the medium irradiance treatment that had the greatest impact on relative telomere lengths, which correlate with long-term fitness (Buracco et al. 2017). UV exposure is known to generate ROS which can damage and shorten telomeres (Heck et al. 2003; Kazerouni et al. 2016; Young 2018), which may explain why relative telomere lengths in animals exposed to medium irradiance UVBR were shorter than relative telomere lengths in animals exposed to the low irradiance treatment. Larvae in the high irradiance UVBR treatment likely experienced the greatest ROS levels, but this effect may have been compensated for by a concomitant increase in molecular defence and repair (including increased antioxidant capacity; Lundsgaard et al. 2020), thus reducing ROS-induced telomere shortening. Such a compensatory response has been observed in western spadefoot toad larvae (*Pelobates cultripes*) exposed to pond drying stress (Buracco et al. 2017), whereby relative telomere lengths remained unaffected, most likely due to an upregulation in antioxidant capacity. Importantly, Burraco et al. (2017) also found that larvae that metamorphosed larger and with more fat reserves had shorter telomeres, reflecting increased metabolic rate, ROS production, and cell division in these animals (Savini et al. 2013; Burraco et al. 2017; Young 2018). A similar effect may have occurred in the present study, whereby the reduced size and energy reserves of animals in the high irradiance UVBR treatment reduced the overall sum of telomeric shortening effects, compared to animals in the medium irradiance UVBR treatment. Our findings support the notion that a trade-off exists between immediate/short-term fitness and long-term survivability and longevity in amphibians responding to environmental stress (Burraco et al. 2017). Further research elucidating the complex interactions between stressors that cause telomere shortening, and variables that counteract telomere shortening (e.g. reduced growth) are warranted.

The finding that an acute, high irradiance exposure to UVBR in the larval period can have short-term and long-term latent effects post-metamorphosis has important implications for our understanding of how this stressor may be shaping amphibian population dynamics. Studies of UVBR exposures in South and Central America suggest a doubling to tripling in the frequency of short-term maximum UVBR exposure events in relatively recent times due to changes in the distribution of ozone and cloud cover (Middleton et al. 2001; Schuch et al. 2015a). It is likely that similar increases in acute UVBR events will continue to occur in some tropical regions due to climate change (Hegglin and Shepherd 2009; Mckenzie et al. 2011; Williamson et al. 2014). Although perturbations to early life-history stages of amphibians may not drive population declines as much as previously assumed (Vonesh and De la Cruz 2002; Vonesh 2005), the present study provides evidence for an additional mechanism by which UVBR exposure in early-life-history stages could shape population dynamics, by way of carryover effects into the crucial juvenile stage. Field measurements taken to estimate the population-scale effects of UVBR exposure must jointly consider the dose and irradiance of exposure to account for the complex interactive effects between these UVBR parameters on long-term amphibian health.

## ACKNOWLEDGEMENTS

The authors would like to thank Isabella Andresen, Katelyn Bell, Ilha Byrne, Jarrod Cameron, Aimee Collier, Douglas Ledlie, Yufei (Lily) Pan, Sophie Smith and Terena Lucas-Thornton for their assistance with animal maintenance, Dr Edward Meyer for spawn collection, and Dr Pat Ward and Dr Simone Blomberg for statistical guidance.

